# TreeN93: a non-parametric distance-based method for inferring viral transmission clusters

**DOI:** 10.1101/383190

**Authors:** Niema Moshiri

## Abstract

**Summary:** Highly-used methods for identifying transmission clusters of rapidly-evolving pathogens from molecular data require a user-determined distance threshold. The choice of threshold is often motivated by epidemiological information known *a priori,* which may be unfeasible for epidemics without rich epidemiological information. TreeN93 is a fully non-parametric distance-based method for transmission cluster identification that scales polynomially.

**Availability and implementation:** TreeN93 is implemented in Python 3 and is freely available at https://github.com/niemasd/TreeN93/.

## 1 Introduction

In the molecular epidemiology of rapidly-evolving pathogens, notably Human Immunodeficiency Virus (HIV), it has become routine to identify clusters of transmission using sequence data(Little *et al.*, 2014; Wertheim *et al.*, 2017). Some methods, such as Cluster Picker (Ragonnet-Cronin *et al.*, 2013) and TreeCluster (Moshiri, 2018a), use the sequences to infer viral phylogenies and subsequently identify clusters from the resulting trees, while other methods, such as HIV-TRACE (Pond *et al.*, 2018), attempt to identify clusters directly from the sequences.

A key limitation of the aforementioned methods is that they are parametric: they require a user-chosen distance threshold *t* with which to cluster individuals. Cluster Picker clusters individuals such that the maximum pairwise *p*-distance of sequences within any cluster is at most *t*, the individuals in a cluster define the leaves of a clade in the viral phylogeny, and the number of clusters is minimized. TreeCluster uses the same objectives as Cluster Picker, but distances are computed directly from the viral phylogeny instead of from sequences. HIV-TRACE clusters individuals such that, if the pairwise TN93 distance (Tamura and Nei, 1993) between the sequences of two individuals is at most t, they are placed in the same cluster, and the number of clusters is maximized. Studies utilizing these methods typically identify clusters using multiple distance thresholds and then validate their results using epidemiological information (Rose *et al.*, 2017), but this form of validation is infeasible when studying epidemics in which rich epidemiological information about the patients is unavailable.

Here, I introduce TreeN93, a threshold-free distance-based method for inferring viral transmission clusters. In addition to identifying transmission clusters from TN93 distances, TreeN93 constructs a data structure that allows for rapid HIV-TRACE cluster identification using any threshold.

## 2 Materials and methods

### 2.1 Implementation

TreeN93 is implemented as a command-line program written in Python 3. It utilizes the TreeSwift package (Moshiri, 2018b) and a built-in Disjoint Set class. The built-in Disjoint Set abstract data type is implemented using an Up-Tree data structure with path compression and union-by-size for amortized 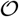(1) time complexity of its find and union operations TreeN93 requires as input the set of all pairwise TN93 distances in the format output by the `tn93` program included in HIV-TRACE.

### 2.2 Clustering objective

Let *d*(*u,v*) denote the distance (e.g. TN93) between sequences *u* and *v*. Given a set of sequences *S* and a number *t* ≥ 0, let *C*(*S|t*) denote a clustering of the sequences of *S* such that, for all pairs of sequences *u,v* ∈ *S*, if *d*(*u,v*) ≤ *t*, *u* and *v* must be in the same cluster, and the number of clusters is maximized. Let *N*(*S|t*) denote the number of clusters in *C*(*S|t*) containing at least two individuals. TreeN93 returns a clustering *C*(*S|t*′) maximizing *N*(*S|t*′) over all *t* ′ ≥ 0. In addition to a clustering, TreeN93 outputs a binary tree *T* such that *C*(*S|t*) can be obtained by cutting *T* at *t* distance above the leaves and creating a cluster containing the leaves of each resulting subtree.

## 3 Results

### 3.1 Time complexity

Given a dataset with *n* individuals (and thus ^(*n*)(*n* – 1)^/_2_ pairwise distances), TreeN93 first sorts the pairwise distances in 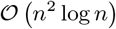, then constructs the TreeN93 tree structure in 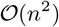, and lastly clusters the individuals in 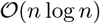, yielding an overall worst-case time complexity of 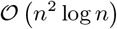 if the input is unsorted and 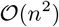 if the input is sorted.

To benchmark execution time as a function of *n*, TreeN93 was executed on 10 replicates of various sizes of *n* on a CentOS 6.6 machine with an Intel Xeon CPU E5-2670 2.60GHz processor (one thread per replicate), and the resulting execution times can be found in Table 1.

**Table 1:**
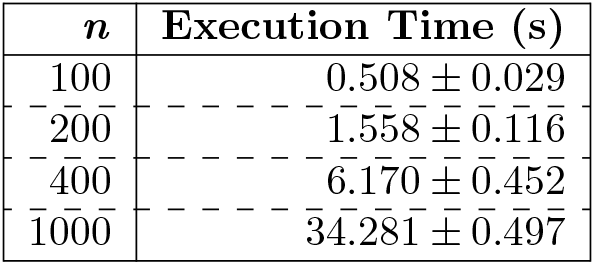
Execution Time

### 3.2 Real-world analysis

To validate TreeN93 as well as to explore its uses, I analyzed three viral sequence datasets: a set of 639 HIV-1 subtype B *pol* sequences (Little *et al.*, 2014), the entire 2016 Los Alamos National Laboratory (LANL) HIV-1 *pol* web alignment (4,416 sequences), and the entire 2008 LANL HCV Core web alignment (783 sequences). For each dataset, I computed the optimal clustering, the corresponding threshold, the number of clusters (i.e., > 1 individual) in the optimal clustering, and the number of singletons (i.e., unclustered individuals) in the optimal clustering. The results can be found in Table 2.

**Table 2:** Real-World Analysis

| Dataset | $n$ | Threshold | Clusters | Singletons |
| --- | --- | --- | --- | --- |
| San Diego HIV | 639 | 0.009991 | 91 | 361 |
| Los Alamos HIV | 4,416 | 0.029034 | 159 | 3,428 |
| Los Alamos HCV | 783 | 0.012341 | 44 | 565 |

Interestingly, using the default HIV-TRACE threshold of 0.015, Little *et al.* (2014) identified 90 transmission clusters in the San Diego HIV-1 dataset, whereas TreeN93 successfully identified 91 transmission clusters in the dataset. Thus, TreeN93 was able to infer transmission clusters with similar performance, but without the need for any prior knowledge regarding appropriate threshold. It must be noted that the default threshold of 0.015 in HIV-1 was largely informed using epidemiological evidence of linkage between partners (Hué *et al.*, 2004), but this information is not typically feasibly obtained for pathogens spread by other means (e.g. foodborne or airborne), emphasizing the importance of a non-parametric method of transmission cluster identification.

## Discussion

As TreeN93 is implemented in Python 3 and is quite light in terms of dependencies, it can be utilized with minimal setup issues on a wide variety of computing platforms. Further, because of its simple code and its command line interface, it can be easily updated or integrated into existing workflows to meet the broad needs of the bioinformatics and epidemiological communities. For example, TreeN93 can parse tn93-format distances from standard input, meaning it can be easily plugged into an existing HIV-TRACE workflow. TreeN93 fills a gap in the current state of transmission cluster identification by providing a fully-nonparametric yet computationally efficient approach.

## Acknowledgements

The author would like to thank Siavash Mirarab for his academic mentorship, and he would like to thank Joel Wertheim, Sergei Pond, Steven Weaver, and Davey Smith for their guidance regarding HIV epidemiology.

## References

Hué, S., Clewley, J. P., Cane, P. A., and Pillay, D. (2004). HIV-1 pol gene variation is sufficient for reconstruction of transmissions in the era of antiretroviral therapy. AIDS (London, England), 18 (5), 719–28.

Little, S. J., Pond, S. L. K., Anderson, C. M., Young, J. A., Wertheim, J. O., Mehta, S. R., May, S., and Smith, D. M. (2014). Using HIV networks to inform real time prevention interventions. PLoS ONE, 9 (6).

Moshiri, N. (2018a). TreeCluster: a Massively scalable transmission clustering using phylogenetic trees. bioRxiv.

Moshiri, N. (2018b). TreeSwift: a massively scalable Python package for trees. bioRxiv.

Pond, S. L. K., Weaver, S., Brown, A. J. L., and Wertheim, J. O. (2018). HIV-TRACE (Transmission Cluster Engine): a tool for large scale molecular epidemiology of HIV-1 and other rapidly evolving pathogens. Molecular Biology and Evolution, (msy016).

Ragonnet-Cronin, M., Hodcroft, E., Hue, S., Fearnhill, E., Delpech, V., Brown, A. J., and Lycett, S. (2013). Automated analysis of phylogenetic clusters. BMC Bioinformatics, 14 (1), 317.

Rose, R., Lamers, S. L., Dollar, J. J., Grabowski, M. K., Hodcroft, E. B., Ragonnet-Cronin, M., Wertheim, J. O., Redd, A. D., German, D., and Laeyendecker, O. (2017). Identifying Transmission Clusters with Cluster Picker and HIV-TRACE. AIDS Research and Human Retroviruses, 33 (3), 211–218.

Tamura, K. and Nei, M. (1993). Estimation of the Number of Nucleotide Substitutions in the Control Region of Mitochondrial DNA in Humans and Chimpanzees ‘. Molecular Biology and Evolution, 10 (3), 512–526.

Wertheim, J., Murrell, B. L., Forgione, L., and Torian (2017). Cluster growth dynamics suggest strategy for targeted intervention in New York City public health HIV-1 surveillance registry. In A. Leigh Brown, A. McLean, E. Hodcroft, J. Albert, M. Kalish, T. Leitner, B. Korber, J. Mullins, S. Kosakovsky Pond, M. Rolland, S. Wolinsky, and M. Worobey, editors, HIV Dynamics & Evolution, number 1122, page 38, Sleat, Isle of Skye, Scotland. UC San Diego School of Medicine.

